# Cdc42 coordinates apical membrane formation and secretory granule maturation through v-SNARE maintenance in salivary acinar cells

**DOI:** 10.1101/2025.07.31.667845

**Authors:** Akiko Shitara, Haruna Nagase, Christopher K. E. Bleck, Yuta Ohno, Hideo Kataoka, Keitaro Satoh, Masanori Kashimata

## Abstract

Epithelial acinar cells of the salivary gland establish apical-basal polarity and secretory competence during development, processes that require tightly coordinated membrane trafficking. The Rho-family GTPase Cdc42 is a key regulator of epithelial polarity, but it has rarely been linked to the vesicle-fusion machinery in vivo. Using mice with acinar cell-specific deletion of Cdc42, we found that Cdc42 loss caused the accumulation of small vesicles near the apical surface, where the water channel AQP5 was retained in clusters and lumen formation was defective. Cdc42-deficient glands showed a selective reduction of two vesicle-associated SNAREs, VAMP2 and VAMP4, whereas VAMP8, the t-SNAREs, and an endocytic marker were unchanged. The corresponding mRNAs were not decreased, indicating that Cdc42 maintains these v-SNAREs through post-transcriptional mechanisms. Cdc42-deficient acinar cells also accumulated condensing vacuoles, a form of immature secretory granule, indicating impaired granule maturation. We propose that Cdc42 coordinates two distinct membrane-remodeling processes by maintaining these v-SNAREs: VAMP2 supports the fusion of AQP5-containing vesicles with the apical membrane, whereas VAMP4 supports secretory granule maturation. These findings identify the vesicle-fusion machinery as a target of Cdc42-dependent polarity control, with potential implications for secretory gland disorders.

## Introduction

Epithelial cells establish tissue integrity through apical-basal polarity. This polarity enables directional transport of proteins and solutes, which is essential for organ function (1, 2). In secretory epithelia such as salivary glands, acinar cells must form polarized structures during development. These structures have distinct apical membrane domains and basolateral membrane domains (3, 4). Proper formation of these polarized structures requires precise coordination of membrane transport pathways. However, the control mechanisms remain poorly understood in vivo.

Small GTPases orchestrate membrane trafficking and epithelial polarity (5, 6). Among them, the Rho-family GTPase Cdc42 is a key upstream regulator: it activates the Par6–aPKC complex, remodels the actin cortex, and guides apical transport in intestine, kidney, and other epithelia (7, 8). During apical membrane biogenesis, Rab11a-positive apical recycling endosomes contribute to cargo delivery to the nascent luminal surface, a process facilitated by Cdc42 (9–11).

Vesicle fusion at the lumen relies on SNARE complexes, formed between vesicle-associated (v-) SNAREs such as the VAMPs and target-membrane (t-) SNAREs such as syntaxins and SNAP proteins (12). In secretory epithelia, secretory vesicles and granules fuse with the luminal membrane via VAMP2- or VAMP8-containing complexes (13–15). Beyond their role at the plasma membrane, v-SNAREs also act earlier in the secretory pathway: VAMP4, together with syntaxin 6, operates at the trans-Golgi network and on immature secretory granules, where it participates in the membrane-remodeling steps that drive granule maturation (16, 17). Thus, distinct v-SNAREs govern different stages of the secretory pathway—from apical membrane fusion to secretory granule maturation. Yet how Cdc42 controls these v-SNARE-mediated processes in vivo remains unknown.

Salivary gland acinar cells develop polarized apical and basolateral domains, together with secretory competence, during late embryonic and early postnatal stages (18). We previously showed that acinar-specific deletion of Cdc42 disrupts apical membrane formation, lumen morphogenesis, and subsequent lumen maintenance (19–21). However, the specific molecular mechanisms remain unclear.

In this study, we demonstrate that Cdc42 orchestrates two membrane-remodeling processes—apical membrane formation and secretory granule maturation—by maintaining the protein levels of two v-SNAREs, VAMP2 and VAMP4, through post-transcriptional mechanisms during salivary gland development.

## Results & Discussion

### Loss of Cdc42 leads to accumulation of vesicles near the apical lumen in salivary gland acinar cells

To explore the role of Cdc42 in membrane organization, we generated acinar cell-specific Cdc42 knockout mice by crossing ACID-Cre, Cdc42^flox/flox^, and mT/mG mice (19, 20). This mouse model was engineered so that Cdc42 knockout is initiated precisely at the onset of acinar cell formation, which begins during embryonic development. Cre recombinase activity was visualized using the mT/mG reporter system (22). In Cdc42-deficient cells, membrane-targeted GFP localized to both the plasma membrane and discrete intracellular vesicles.

These GFP-positive vesicles did not exhibit a random cytoplasmic distribution; instead, they were enriched in regions adjacent to the apical surface (Figure 1A). Three-dimensional reconstruction confirmed that approximately 50% of the GFP clusters were located within 1 µm of the apical lumen (Figure 1B). Transmission electron microscopy showed numerous small vesicles (approximately 0.1–0.25 µm) clustered near the luminal membrane (Figure 1C), which were largely absent from Cdc42 control glands. Quantification confirmed this accumulation: the proportion of small vesicles (≤0.3 µm², likely including endosomes and transport vesicles) among total vesicles was significantly increased in Cdc42-deficient acini (Figure 1D).

**Figure 1.**
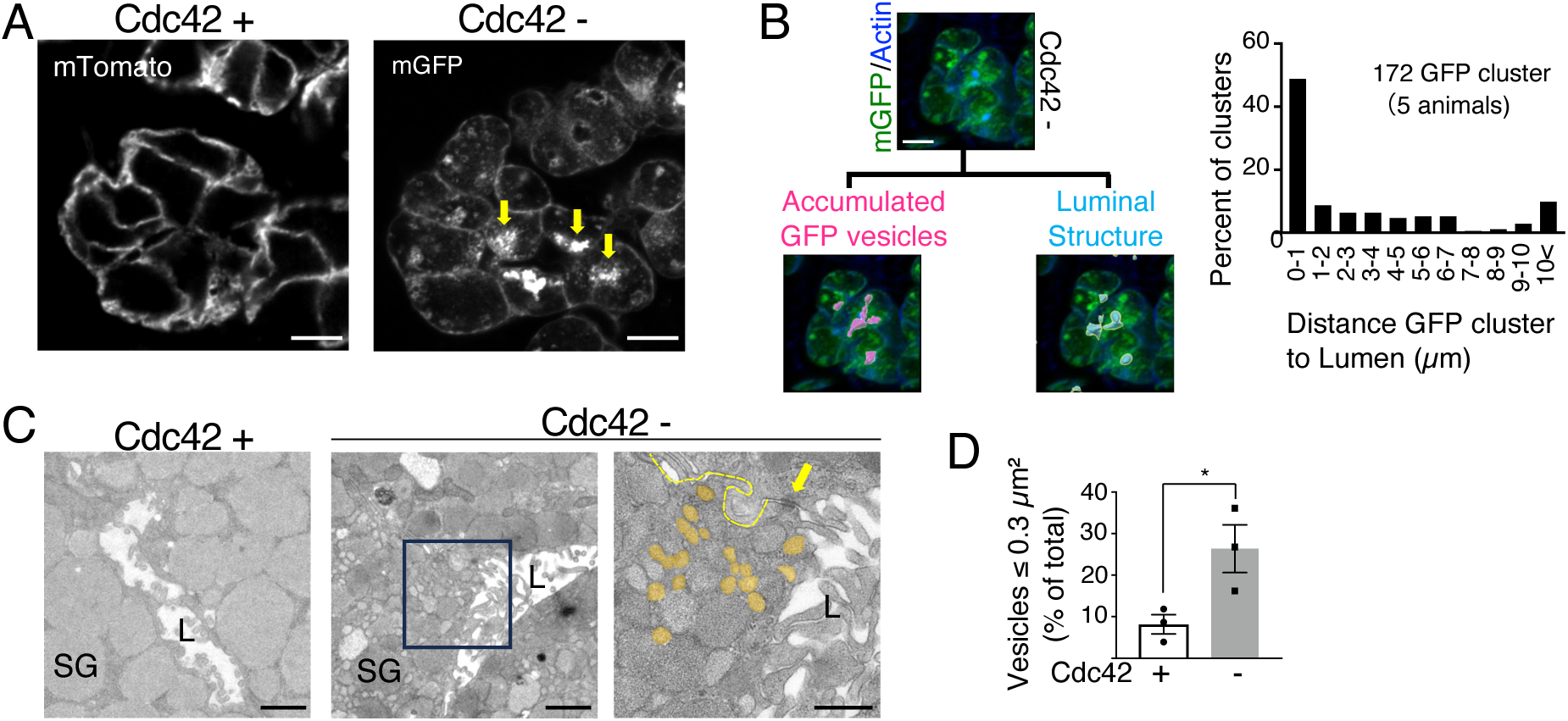
Cdc42 deletion causes accumulation of small vesicles beneath the apical surface. (A) Representative confocal images of parotid gland tissue from control (Cdc42□) and acinar-specific Cdc42 knockout (Cdc42□) mice. Plasma membranes are labeled by mTomato (left) or Cre-activated mGFP (right); yellow arrows indicate mGFP-positive vesicle clusters. Scale bar, 10 µm. (B) Spatial analysis of vesicle-to-lumen distance. Left, workflow: lumina were visualized with phalloidin-405 (blue), and mGFP clusters (magenta) and luminal surfaces (cyan) were rendered in Imaris to calculate shortest distances. Right, histogram of 172 mGFP clusters, predominantly located within 1 µm of the lumen (five Cdc42□ mice). Scale bar, 10 µm. (C) Transmission electron micrographs of acinar cells. Left, control gland (Cdc42□). Middle, Cdc42 gland showing vesicle accumulation near the lumen. Right, enlargement of the boxed region; vesicles ≤0.3 µm² are pseudo-colored orange, cell boundaries are traced by yellow dashed lines, and a tight junction is marked by a yellow arrow. SG, secretory granule; L, lumen. Scale bars, 1 µm (left, middle) and 500 nm (right). (D) Quantification of the proportion of small vesicles (≤0.3 µm², likely including endosomes and transport vesicles) relative to total vesicles. Cdc42□, 8.2 ± 4.0%; Cdc42□, 26.4 ± 10.0% (mean ± S.D.; n = 3 mice per genotype; P = 0.042, unpaired t-test). Orange-colored vesicles in (C) correspond to the small vesicles quantified here.

The accumulation of vesicles near the apical lumen mirrors earlier 3D-culture findings in which Cdc42 loss disrupted apical membrane identity and vesicle targeting (9, 23). Importantly, our tissue-based observations reveal that these GFP-positive vesicle clusters are located within approximately 1 µm of the apical surface. This spatial constraint suggests that Cdc42 acts primarily in the final stages of vesicle–membrane fusion rather than in long-range trafficking events.

### Cdc42 deficiency perturbs the localization of AQP5, Rab11a, and NKCC1 in distinct ways

To determine the identity and behavior of the proteins accumulating near the apical lumen in Cdc42-deficient acinar cells, we examined AQP5, a water channel predominantly localized to the apical plasma membrane (18, 24); Rab11a, a small GTPase that labels apical recycling endosomes essential for apical membrane biogenesis (10); and the basolateral transporter NKCC1 (25).

In Cdc42 control acinar cells, AQP5 showed its expected apical and basolateral distribution, Rab11a was present weakly in the cytoplasm and more prominently near the basolateral membrane, and NKCC1 was confined to the basolateral membrane (Figure 2A–C). In Cdc42-deficient cells, all three proteins showed increased cytoplasmic fluorescence intensity (Figure 2D–F), indicating that Cdc42 loss leads to intracellular accumulation of cargoes normally destined for distinct membrane domains.

**Figure 2.**
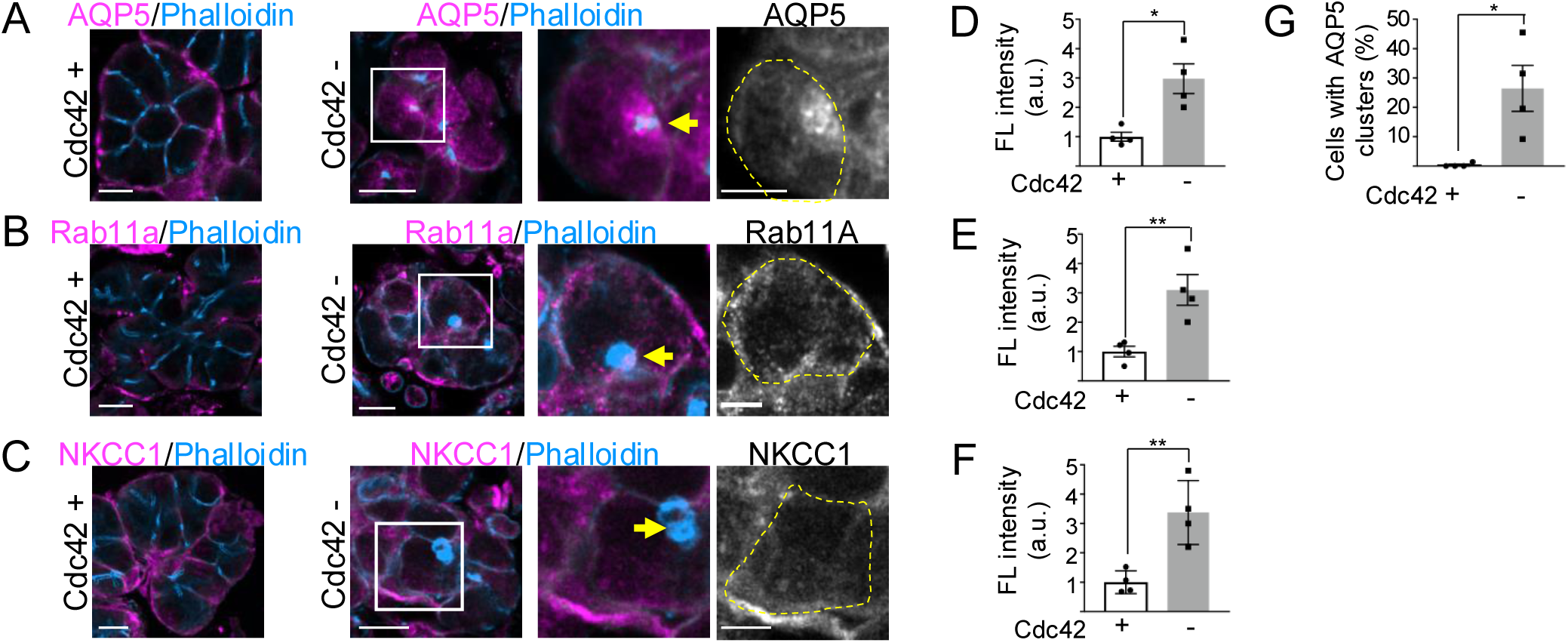
Cdc42 deficiency perturbs the localization of AQP5, Rab11a, and NKCC1 in distinct ways. (A–C) Immunostaining for AQP5 (A), Rab11a (B), and NKCC1 (C) (magenta) together with phalloidin (cyan) in control (Cdc42□) and Cdc42□ acinar cells. For each protein, the left panels show low-magnification views of control and Cdc42□ glands; boxed regions in the Cdc42□ panels are shown as higher-magnification views, and grayscale panels show the single (target) channel with the cell outline (yellow dashed line). Yellow arrows indicate the apical surface. Scale bars, 10 µm (low magnification) and 5 µm (higher-magnification views). (D–F) Quantification of cytoplasmic fluorescence intensity of AQP5 (D), Rab11a (E), and NKCC1 (F), measured as the mean intensity within a fixed-area region of the subapical cytoplasm, normalized to the control mean (mean ± S.D.). AQP5, 3.0-fold increase (P = 0.0098; n = 4 mice per genotype); Rab11a, 3.1-fold increase (P = 0.0089; n = 4 mice per genotype); NKCC1, 3.4-fold increase (P = 0.006; n = 4 mice per genotype). *P < 0.05, **P < 0.01 by unpaired t-test. (G) Proportion of acinar cells containing peri-membrane AQP5 clusters. Cdc42□, 26.5 ± 15.6%; Cdc42□, 0.35 ± 0.70% (mean ± S.D.; P = 0.0265, Mann–Whitney U test; n = 4 mice per genotype). Cluster quantification was performed only for AQP5, which formed discrete clusters. The same criterion applied to Rab11a yielded essentially no cluster-positive signal; Rab11a and NKCC1 showed diffuse cytoplasmic increases and were quantified by intensity. See Supplementary Table S1 for antibody details.

Importantly, the pattern of accumulation differed among these proteins. AQP5 formed discrete clusters near the apical membrane; quantification showed that a substantial fraction of Cdc42-deficient acinar cells contained such AQP5 clusters, whereas these were essentially absent in controls (Figure 2G). In contrast, when assessed by the same criterion, Rab11a did not form comparable discrete clusters; instead, its signal increased both in the cytoplasm and near the basolateral membrane, indicating a diffuse, overall increase rather than the focal clustering seen for AQP5. NKCC1 also showed a diffuse increase: its basolateral localization was retained, while its signal became detectable throughout the cytoplasm.

These results indicate that Cdc42 deficiency perturbs the trafficking of multiple cargoes, but in distinct ways. The most prominent phenotype is the clustered retention of AQP5 near the apical membrane, consistent with a failure of apical cargo to be delivered to or fused with the apical surface. The diffuse redistribution of Rab11a, distinct from the AQP5 clusters, suggests a broader perturbation of the recycling/endosomal system rather than the trapping of a single discrete compartment. The intracellular appearance of NKCC1 indicates that basolateral trafficking is also affected, though without the clustered retention seen for AQP5. Thus, rather than a strictly apical-selective defect, Cdc42 loss affects multiple trafficking routes, with the clustered retention of apical AQP5 as its most striking feature.

### Cdc42 deficiency selectively reduces the v-SNAREs VAMP2 and VAMP4 at the post-transcriptional level

The accumulation of cargo-laden vesicles near the apical surface could result from either impaired fusion with the apical membrane or excessive endocytosis. To distinguish between these possibilities, we examined the vesicle fusion machinery and endocytic markers by Western blotting.

Among v-SNAREs, VAMP2 and VAMP4 were both markedly reduced in Cdc42-deficient parotid glands—by approximately 55% and 59%, respectively, relative to α-tubulin—whereas VAMP8 was unchanged (Figure 3A, B). The comparable reduction of two distinct v-SNAREs, VAMP2 and VAMP4, suggested a coordinated effect on a specific subset of the fusion machinery. To define this specificity further, we examined the t-SNAREs SNAP23 and Syntaxin3, which mediate apical membrane fusion, and Syntaxin6, which functions in immature granule fusion during maturation (16). None of these t-SNAREs was reduced: SNAP23 and Syntaxin6 were unchanged, and Syntaxin3 showed a non-significant trend toward increase (Supplementary Figure S1). The endocytic tethering factor EEA1 was also unchanged (Supplementary Figure S1). This dissociation—reduced v-SNAREs with preserved t-SNAREs and endocytic machinery—indicates that Cdc42 deficiency does not globally compromise the fusion apparatus or promote excessive endocytosis, but selectively lowers a specific subset of v-SNAREs.

**Figure 3.**
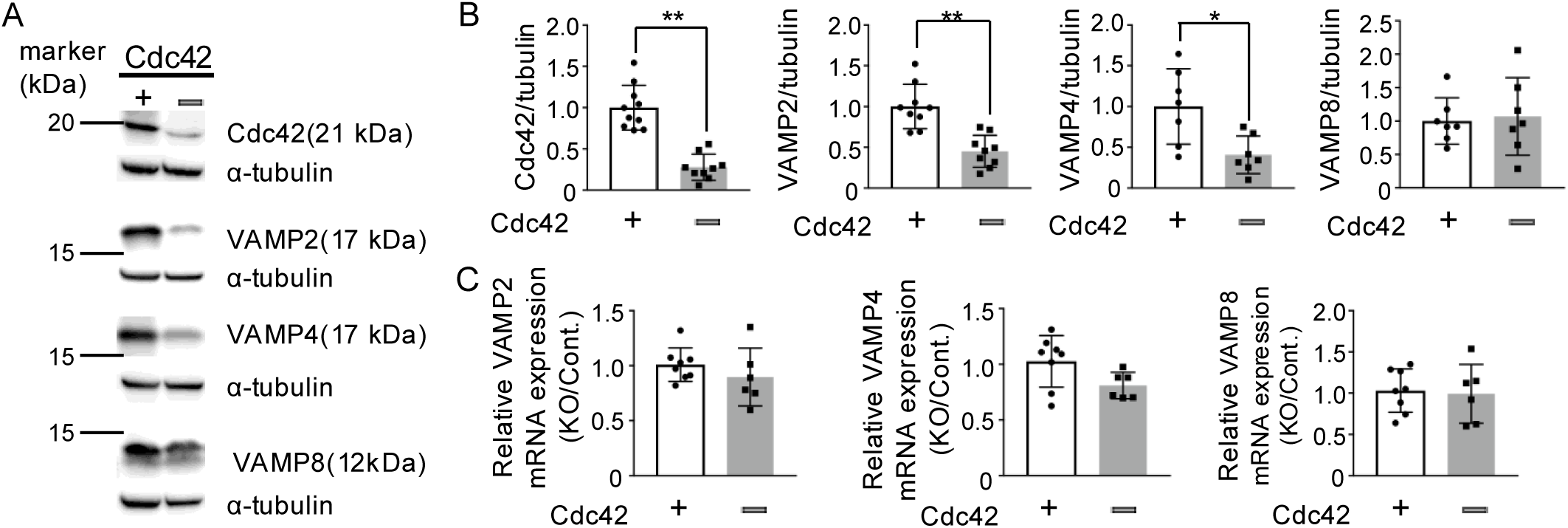
Cdc42 deficiency selectively reduces the v-SNAREs VAMP2 and VAMP4 at the post-transcriptional level. (A) Western blot analysis of the indicated proteins in parotid glands from control (Cdc42□) and Cdc42 mice. Representative blots; molecular weights are indicated. (B) Quantification normalized to α-tubulin (mean ± S.D.). VAMP2 (55% reduction, P = 0.0002; n = 9 per group), VAMP4 (59% reduction, P = 0.0104; n = 7 per group), and VAMP8 (unchanged, P = 0.78; n = 7 per group). Cdc42 (P < 0.0001; n = 10 control, 9 knockout) confirms knockout. *P < 0.05, **P < 0.01 by unpaired t-test. (C) qRT-PCR of Vamp2, Vamp4, and Vamp8 mRNA, normalized to the geometric mean of Rplp0 and Ppia (mean ± S.D.). Vamp2 (P = 0.33; n = 8 control, 6 knockout), Vamp4 (P = 0.058, not significant; n = 8 control, 6 knockout), Vamp8 (unchanged; n = 8 control, 6 knockout).

To determine whether these reductions occurred at the transcriptional or post-transcriptional level, we performed quantitative RT-PCR. Vamp2 mRNA was unchanged despite the marked decrease in VAMP2 protein, and Vamp8 mRNA was likewise unchanged (Figure 3C). For Vamp4, mRNA was reduced by only ∼21% (P = 0.059, not significant; n = 8 control, 6 knockout); a transcriptional change of this magnitude cannot account for the ∼59% loss of VAMP4 protein (Figure 3C). Together, these results indicate that Cdc42 maintains VAMP2 and VAMP4 predominantly through post-transcriptional mechanisms.

The trend toward increased Syntaxin3, although not statistically significant, may reflect a compensatory response to the reduction of its cognate v-SNAREs, although this remains to be tested. Reduced VAMP2 provides a molecular explanation for the vesicle accumulation phenotype. With less VAMP2, apical cargo is transported to the apical region but fails to fuse with the luminal membrane, accumulating near the apical surface. VAMP4, by contrast, acts in secretory granule maturation, which we examine next.

### Cdc42 deficiency causes accumulation of condensing vacuoles, indicating impaired secretory granule maturation

VAMP4 operates at the trans-Golgi network and on immature secretory granules, where it is sorted away during granule maturation (16, 17). Its reduction therefore prompted us to examine secretory granule maturation. Our previous study, using the same acinar-specific Cdc42 knockout model, noted heterogeneous electron-dense areas in secretory granules consistent with condensing vacuoles, but that observation was qualitative and was not quantified (19).

Here, in independent micrographs from this model, we quantified condensing vacuoles—immature secretory granules characterized by a partially condensed, electron-dense core surrounded by less electron-dense material. The proportion of secretory granules displaying condensing-vacuole morphology was markedly increased in Cdc42-deficient acini compared with controls (Figure 4). Because condensing vacuoles are a recognized form of immature secretory granule (26, 27), their accumulation indicates a block in granule maturation.

**Figure 4.**
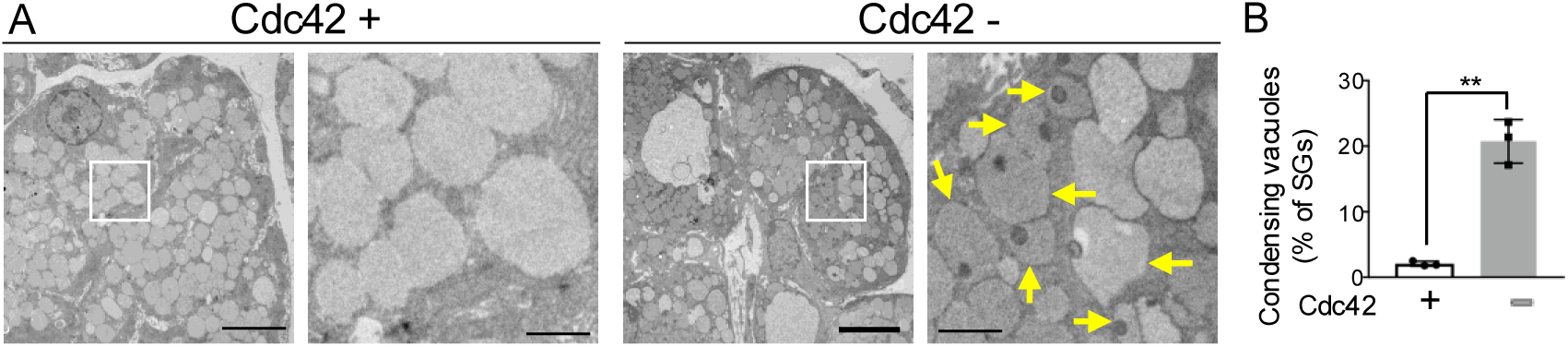
Cdc42 deficiency causes accumulation of condensing vacuoles, indicating impaired secretory granule maturation. (A) Transmission electron micrographs of acinar cells from control (Cdc42□, left) and Cdc42□ (right) glands, shown at low magnification (left of each pair) and high magnification (right of each pair). In control glands, secretory granules were uniformly electron-dense, whereas in Cdc42□ glands, granules with condensing-vacuole morphology—a partially condensed, electron-dense core surrounded by less electron-dense material—were frequently observed (yellow arrows). Scale bars, 5 µm (low magnification) and 1 µm (high magnification). (B) Proportion of secretory granules displaying condensing-vacuole morphology (Cdc42□, 2.1 ± 0.4%; Cdc42□, 20.7 ± 3.3%; mean ± S.D.; n = 3 mice per genotype; P = 0.0096, Welch’s t-test). The micrographs are from the same acinar-specific Cdc42 knockout model as in our previous study (19) but are not the images shown there.

The concurrent, post-transcriptional loss of VAMP4 provides a candidate molecular basis for this defect. VAMP4 participates in the membrane-remodeling steps that accompany granule maturation, and its reduction would be expected to impair the transition from immature to mature granules, linking our earlier morphological observation (19) to the loss of VAMP4.

### Cdc42 coordinates apical membrane formation and secretory granule maturation through maintenance of VAMP2 and VAMP4

Our findings support a model in which Cdc42 maintains the protein levels of two v-SNAREs, VAMP2 and VAMP4, through post-transcriptional mechanisms, thereby coordinating two distinct membrane-remodeling processes during acinar cell development (Figure 5). Maintenance of VAMP2 supports fusion at the apical membrane; its loss leaves apical cargo, including AQP5, retained in clusters near the apical surface, with defective lumen formation. Maintenance of VAMP4 supports secretory granule maturation; its loss leads to the accumulation of condensing vacuoles. In this way, a single upstream regulator sets the abundance of these two v-SNAREs to govern two membrane processes.

**Figure 5.**
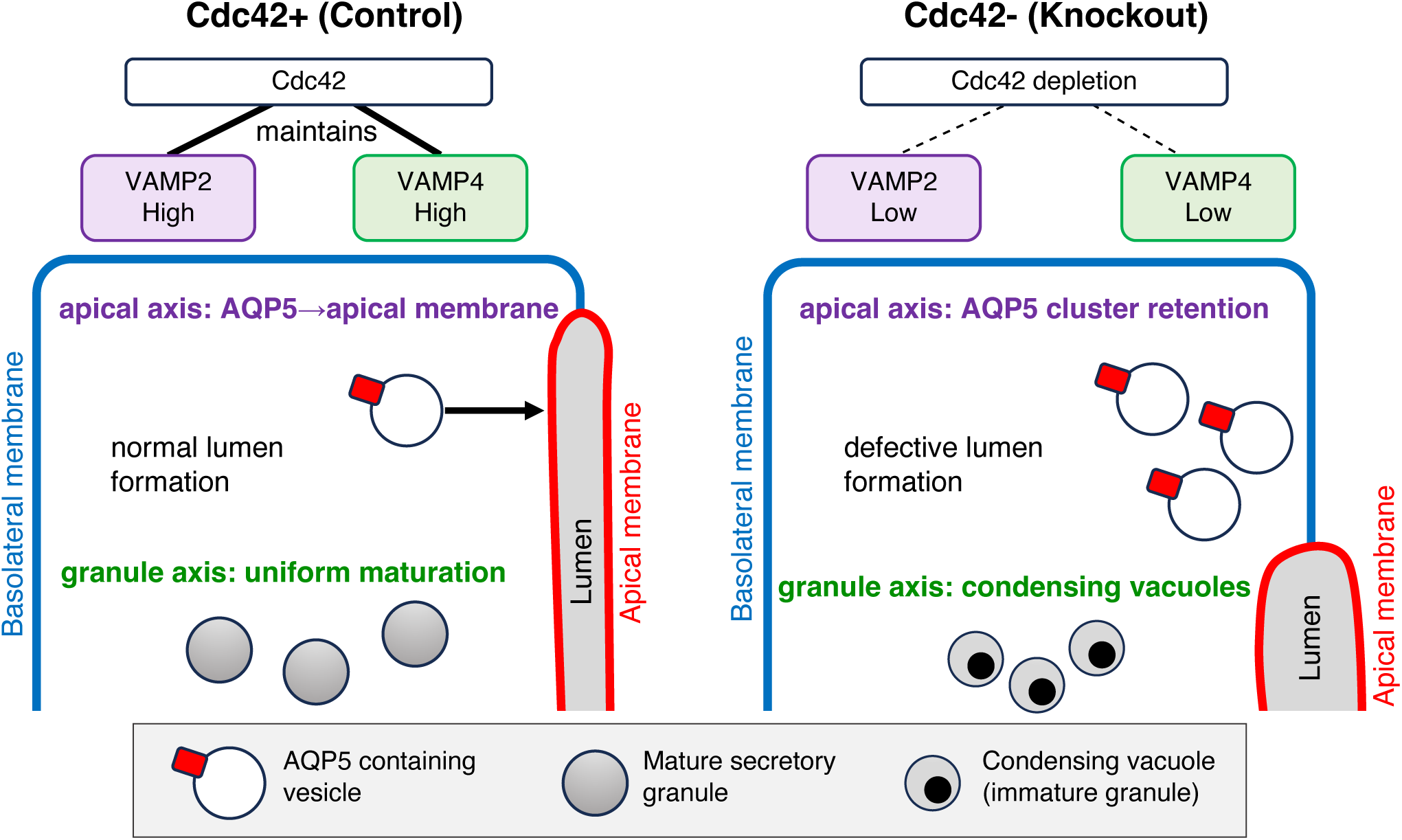
Proposed model: Cdc42 maintains VAMP2 and VAMP4 to coordinate apical membrane formation and secretory granule maturation. In control acinar cells (left), Cdc42 maintains the protein levels of two v-SNAREs, VAMP2 and VAMP4. Along the apical axis, VAMP2 supports fusion of AQP5-containing vesicles to the apical membrane, allowing normal lumen formation. Along the granule axis, VAMP4 supports uniform maturation of secretory granules. In Cdc42-deficient cells (right), reduced VAMP2 and VAMP4 lead to retention of AQP5-containing vesicles in clusters near the apical surface with defective lumen formation, and to accumulation of condensing vacuoles (immature granules). Symbols: red square, AQP5; open circle, AQP5-containing vesicle; gray filled circle, mature secretory granule; circle with dense core, condensing vacuole; blue line, basolateral membrane; red line, apical membrane. The shorter lumen in the knockout depicts defective lumen formation.

The stability of VAMP8—which mediates regulated exocytosis of mature granules—is consistent with our earlier observation that stimulus-induced fusion of secretory granules still occurs in this model, albeit with delayed integration into the apical membrane (19, 20). The phenotype we describe therefore reflects a specific impairment of VAMP2- and VAMP4-dependent processes during development.

The mechanism by which Cdc42 sustains these v-SNARE levels remains to be defined, but a direct interaction offers a plausible framework. In pancreatic β cells, Cdc42 binds VAMP2 directly, independently of its nucleotide state (28). Loss of Cdc42 protein might therefore destabilize VAMP2, and possibly VAMP4, leading to their post-transcriptional reduction. To our knowledge, a link between Cdc42 and VAMP4 has not been reported previously, and its association with impaired granule maturation identifies a new candidate mechanism that warrants further investigation. We also examined whether major degradation pathways could account for the selective v-SNARE loss. Global ubiquitination and p62 levels were unchanged, and although the LC3B-II/LC3B-I ratio was elevated, p62 did not accumulate. This pattern is not readily explained by a simple block in autophagic flux, although increased autophagic flux cannot be excluded without direct flux measurements (e.g., in the presence of bafilomycin A1) (Supplementary Figure S2). These observations argue against a global activation of the ubiquitin–proteasome system or a general autophagic defect as the basis for the selective reduction of VAMP2 and VAMP4, though the specific degradative or translational mechanism remains to be identified.

Definitive causal testing—for example, v-SNARE re-expression or measurement of protein half-life—would ideally be performed in cultured cells. However, salivary gland acinar cells rapidly dedifferentiate and lose their secretory phenotype and granules in conventional culture (29, 30), and no established system reproduces the apical membrane formation and granule maturation observed in vivo in our developmental model; recent three-dimensional and organoid approaches can partially restore epithelial polarity but are largely derived from stem/progenitor cells and remain under optimization (31, 32). We therefore relied on in vivo analysis in polarized tissue, which retains the physiological context lost in dedifferentiated cultures.

In conclusion, this study identifies Cdc42 as a coordinator of apical membrane formation and secretory granule maturation during acinar cell development, acting through post-transcriptional maintenance of the v-SNAREs VAMP2 and VAMP4. These findings reveal how a polarity regulator can integrate distinct membrane-trafficking processes through control of the fusion machinery, and may have implications for epithelial disorders in secretory tissues. Whether this mechanism is conserved across secretory epithelia, and whether its disruption contributes to glandular disease, will be important questions for future study.

## Experimental Procedures

### 1. Mouse Models and Ethical Approval

All animal procedures were approved by the Animal Experiment Committee of Asahi University (approval numbers 22-011, 23-025, 24-029, 25-032) and conducted in accordance with institutional guidelines.

mT/mG reporter mice were obtained from The Jackson Laboratory (Bar Harbor, ME, USA). Cdc42^flox/flox^ mice and ACID-Cre (AQP5-Cre) mice were kindly provided by Dr. Y. Zheng (Cincinnati Children’s Hospital Medical Center, OH, USA)(33) and Dr. Z. Borok (University of Southern California, CA, USA)(34), respectively. To generate acinar cell-specific Cdc42-deficient mice, Cdc42^flox/flox^; AQP5-Cre; mT/mG mice were generated by crossing the respective lines as described previously (19, 20). Both male and female mice aged 10–35 weeks and weighing 20–40 g were used. Data from both sexes were pooled for analysis. Because the number of animals per group was not powered to detect sex differences, sex was not included as a variable in the statistical analyses; no overt sex-dependent differences in the phenotypes were noted. Animals were housed under standard conditions (12 h light/dark cycle, 22 ± 2 °C, 50 ± 10% humidity) with ad libitum access to food and water.

### 2. Tissue Collection and Processing

Mice were anesthetized by intraperitoneal injection of ketamine (100 mg/kg) and xylazine (20 mg/kg), followed by transcardial perfusion with 4% paraformaldehyde (PFA) in phosphate-buffered saline (PBS). Parotid glands were harvested and post-fixed overnight in 4% PFA at 4 °C. Tissues were cryoprotected in 15% and 30% sucrose in PBS, embedded in OCT compound (Sakura Finetek, Torrance, CA, USA), and frozen in 2-methylbutane cooled with dry ice. Cryosections (10 µm) were prepared using a CM1850 cryostat (Leica Biosystems, Wetzlar, Germany) for immunostaining.

### 3. Immunofluorescence Staining and Confocal Imaging

Sections were permeabilized and blocked in PBS containing 0.02% saponin and 10% fetal bovine serum (FBS). Primary antibodies were applied overnight at 4 °C. The following day, Alexa Fluor-conjugated secondary antibodies were applied for 45 min at room temperature. Phalloidin-405 was used for F-actin visualization when applicable. Details of all antibodies and reagents are listed in Supplementary Table S1. Slides were mounted with Fluoromount (Diagnostic BioSystems, Pleasanton, CA, USA) and imaged using an LSM980 confocal microscope (Carl Zeiss, Oberkochen, Germany) with Zen software. For comparative analysis, control (Cdc42□) and Cdc42-deficient samples were imaged under identical acquisition parameters, and images were displayed with the same brightness and contrast settings. Images were analyzed using Imaris (Oxford Instruments, Abingdon, UK) and Fiji (ImageJ, NIH, Bethesda, MD, USA).

### 4. Transmission Electron Microscopy

Salivary glands were fixed for 90 min in 0.1 M phosphate buffer (pH 7.2) containing 2% glutaraldehyde and 2% PFA (Electron Microscopy Sciences, Hatfield, PA, USA). Post-fixation with aqueous 1% osmium tetroxide and en bloc staining with 1% uranyl acetate were performed. Samples were dehydrated in graded ethanol and embedded in EMbed-812 resin (Electron Microscopy Sciences). Ultrathin sections were stained with uranyl acetate and lead citrate and examined with a JEM-1200EX transmission electron microscope (JEOL, Tokyo, Japan) at 80 kV. Micrographs were captured with a bottom-mounted 6-megapixel CCD camera (Advanced Microscopy Techniques, Woburn, MA, USA).

### 5. Western Blotting

Parotid glands were dissected from anesthetized mice, homogenized in RIPA buffer containing protease/phosphatase inhibitors and 0.5 mM PMSF, and incubated on ice for 15 min. For the ubiquitination (FK2) analysis, 20 mM N-ethylmaleimide (NEM) was additionally included in the RIPA buffer to inhibit deubiquitinating enzymes and preserve ubiquitin conjugates. Lysates were centrifuged at 14,000×g (4 °C, 10 min), and supernatants were collected. Protein concentration was determined via the Bradford assay. Equal amounts (20 µg) were resolved by SDS-PAGE and transferred to PVDF membranes (Trans-Blot Turbo, Bio-Rad, Hercules, CA, USA).

Immunoblotting was performed by one of two methods, as indicated for each antibody in Supplementary Table S1. For the conventional method, membranes were blocked in 10% skim milk in TBS-T at 4 °C overnight, followed by antibody incubation using an enhanced rapid immunoblotting protocol as described previously (35, 36); primary antibodies were diluted in Can Get Signal® Solution 1 (Toyobo, Osaka, Japan) and HRP-conjugated secondary antibodies in Can Get Signal® Solution 2. For the remaining antibodies (see Supplementary Table S1), immunoblotting was performed using the iBind Automated Western System (Thermo Fisher Scientific, Waltham, MA, USA) according to the manufacturer’s instructions. Protein bands were visualized using enhanced chemiluminescence (ECL) reagent (Cytiva, Marlborough, MA, USA). Chemiluminescence signals were captured with a LuminoGraph III imaging system (ATTO, Tokyo, Japan). Band intensities were quantified using Fiji and normalized to α-tubulin. Both detection methods gave comparable results, and each target protein was normalized to α-tubulin detected by the same method. For ubiquitinated proteins (FK2), the total signal across the 30–250 kDa range of each lane was quantified and normalized to α-tubulin. Each target protein was detected on a separate membrane using an independent set of gland lysates and normalized to α-tubulin on the same membrane; consequently, the number of biological replicates (n) differs among targets, with exact values given in the figure legends. See Supplementary Table S1 for antibody details.

### 6. Quantitative RT-PCR

Total RNA was extracted from parotid glands using the FavorPrep Tissue Total RNA Mini Kit (Favorgen, Pingtung, Taiwan) with DNase I treatment. cDNA was synthesized from 200 ng RNA using ReverTra Ace qPCR RT Master Mix (Toyobo, Osaka, Japan). qPCR was performed using SYBR Green on a StepOnePlus system (Applied Biosystems, Foster City, CA, USA). Expression was normalized to the geometric mean of Rplp0 and Ppia using the ΔΔCt method.

Primer sequences were: VAMP2 (F: 5′-GCTGGATGACCGTGCAGAT-3′, R: 5′-GATGGCGCAGATCACTCCC-3′); VAMP4 (F: 5’- TTCAAGCGCCACCTAAATGAT-3’, R: 5’- CCAGATGGTCCCCGTAGAAAAA-3’); VAMP8 (F: 5′-AGTGGGAGTGCCGGAAATG-3′, R: 5′-TGAAGTGTTCAGACGTGGCTT-3′); Rplp0 (F: 5′-GGACCCGAGAAGACCTCCTT-3′, R: 5′-GCACATCACTCAGAATTTCAATGG-3′); Ppia (F: 5′-GCACTGCCAAGACTGAATGG-3′, R: 5′-TGCCTTCTTTCACCTTCCCAA-3′).

### 7. Image Quantification

#### Distance from GFP clusters to lumen (Fig. 1B)

Imaris software was used to generate 3D surface models for GFP-positive clusters and lumens. The distance between clusters and lumen surfaces was computed using the “shortest distance calculation” function. Thresholds were determined visually.

#### Vesicle and secretory granule quantification by TEM (Fig. 1D, Fig. 4)

Vesicles and secretory granules were traced manually in Fiji from the same set of electron micrographs. The proportion of small vesicles (≤0.3 µm²) among total vesicles (Fig. 1D) and the proportion of secretory granules showing condensing-vacuole morphology—defined as a partially condensed, electron-dense core surrounded by less electron-dense material (Fig. 4)—were each determined per animal (n = 3 mice per genotype; 3–7 fields and 121–897 vesicles/granules analyzed per animal).

#### Fluorescence intensity and AQP5 cluster quantification (Fig. 2D–G)

Confocal images of control and Cdc42-deficient glands were acquired under identical settings as z-stacks. For each acinar structure, 7 consecutive optical sections in which the acinar lumen was clearly visible were selected from the 13 acquired sections and combined into a single image by sum-slice z-projection in Fiji. In the projected image, a circular region of interest (ROI; area 9.72 µm²) was placed in the subapical cytoplasm of each acinar cell, and the mean fluorescence intensity of the target protein (AQP5, Rab11a, or NKCC1) within the ROI was measured. For cytoplasmic intensity (Fig. 2D–F), these values were averaged per animal and normalized to the control mean (n = 4 mice per genotype, three fields each). The same ROI measurements were used for AQP5 cluster quantification (below). The same procedure was applied to control and Cdc42-deficient samples.

#### AQP5 cluster quantification (Fig. 2G)

For AQP5 cluster quantification, the same ROI measurements described above were used: cells with a mean ROI intensity ≥25,000 (arbitrary units) were scored as AQP5 cluster-positive. This threshold cleanly separated the two genotypes: essentially no control cells reached this value (0.35% cluster-positive), whereas cluster-positive cells were frequent in Cdc42-deficient glands. The percentage of cluster-positive cells was determined per animal (n = 4 mice per genotype, three fields each). This cluster criterion was designed to capture the discrete peri-membrane accumulation characteristic of AQP5. The same criterion was also applied to Rab11a, which showed essentially no cluster-positive signal at the peri-membrane region. Cluster-positive percentages are therefore reported for AQP5 (Fig. 2G), whereas Rab11a and NKCC1, which showed diffuse cytoplasmic increases, were evaluated by cytoplasmic intensity (Fig. 2E, F).

### 8. Exclusion Criteria

For the Cdc42 immunoblot (Figure 3B), a subset of lanes represented non-independent samples derived from the same animal; in each such case, one lane per animal was excluded from quantification so that only one value per animal contributed to the analysis. The excluded lanes are indicated in the uncropped blots (Supporting Information). All other targets were analyzed using independent samples, one per animal. In addition, for the ubiquitination (FK2) analysis (Supplementary Figure S2C, D), one control sample showed markedly reduced α-tubulin signal relative to all other samples, indicating probable loading failure or protein degradation, and was excluded from quantification (final n = 3 control, 4 knockout). The excluded lane is shown in the uncropped blot (Supporting Information).

### 9. Statistical Analysis

All data are presented as mean ± S.D. with individual data points superimposed. Comparisons between two groups were performed using the two-tailed unpaired Student’s t-test; equality of variances was assessed by an F-test, and Welch’s correction was applied when variances differed significantly (F-test P < 0.05). For the proportion of AQP5 cluster-positive cells, where one group had near-zero variance, the Mann–Whitney U test was used. Statistical significance was defined as P < 0.05, and exact P values are reported in the figure legends. All analyses were performed using GraphPad Prism 7 (GraphPad Software, San Diego, CA, USA; www.graphpad.com).

## Data availability

The data supporting the findings of this study are available within the article and its Supporting Information, which includes uncropped Western blot images. Additional raw data (e.g., source values for quantification and qRT-PCR) are available from the corresponding author upon reasonable request.

## Acknowledgments

We thank Ms Riyako Terazawa and Ms Chikako Katayama for expert technical assistance. We are grateful to Dr Roberto Weigert (National Institutes of Health) for sharing the compound *Aqp5-Cre; Cdc42*^flox/flox^ mouse line, which was generated in his laboratory from the *Aqp5-Cre* mice kindly provided by Drs Zea Borok, Edward Crandall, and Peter Flodby (University of Southern California) and the *Cdc42^f^*^lox/flox^ mice generously provided by Dr Yi Zheng (Cincinnati Children’s Hospital Medical Center). We also thank Ms Heba Mohammed for her assistance with electron-microscopy sample preparation. Confocal images were acquired at the Optics and Imaging Facility, NIBB Trans-Scale Biology Center.

This work was supported by JSPS KAKENHI grants 19K20764, 21K10126, 24K20125, 25K13075, and 25K13300, by the Miyata Research Encouragement Grant A (grant no. A24014, Asahi University, JP), by the Koshiyama Research Grant, and by the Ogawa Science and Technology Foundation.

