## Supplementary figures and images for "Cdc42 coordinates apical membrane formation and secretory granule maturation through v-SNARE maintenance in salivary acinar cells"

### Fig.S1

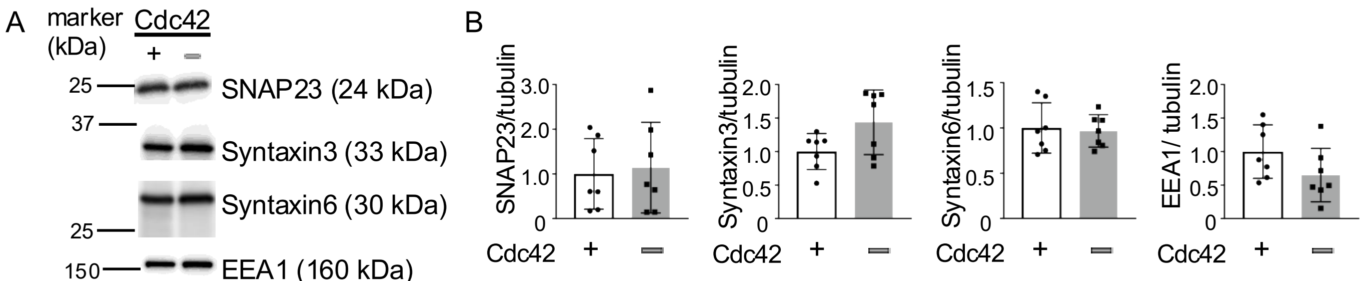

### Fig.S2

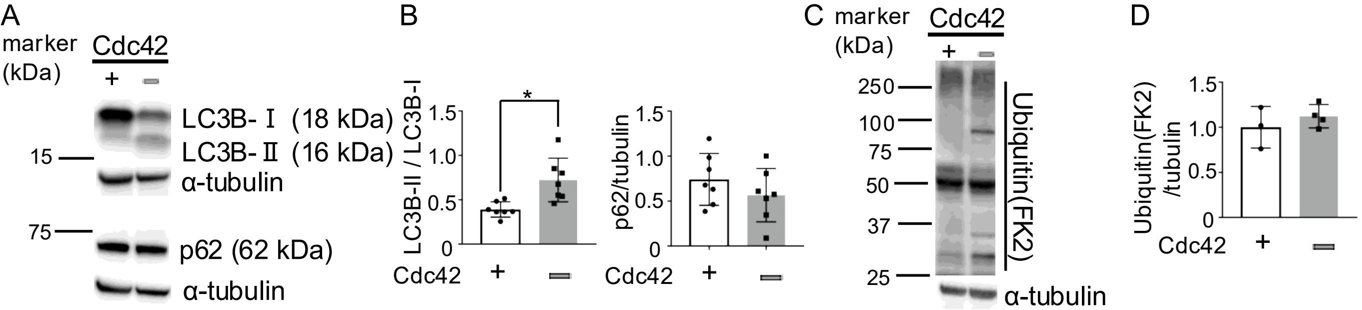
